# Past–future information bottleneck framework for simultaneously sampling biomolecular reaction coordinate, thermodynamics and kinetics

**DOI:** 10.1101/507822

**Authors:** Yihang Wang, João Marcelo Lamim Ribeiro, Pratyush Tiwary

## Abstract

The ability to rapidly learn from high-dimensional data to make reliable bets about the future outcomes is crucial in many contexts. This could be a fly avoiding predators, or the retina processing gigabytes of data almost instantaneously to guide complex human actions. In this work we draw parallels between such tasks, and the efficient sampling of complex biomolecules with hundreds of thousands of atoms. For this we use the Predictive Information Bottleneck (PIB) framework developed and used for the first two classes of problems, and re-formulate it for the sampling of biomolecular structure and dynamics, especially when plagued with rare events. Our method considers a given biomolecular trajectory expressed in terms of order parameters or basis functions, and uses a deep neural network to learn the minimally complex yet most predictive aspects of this trajectory, viz the PIB. This information is used to perform iterative rounds of biased simulations that enhance the sampling along the PIB to gradually improve its accuracy, directly obtaining associated thermodynamic and kinetic information. We demonstrate the method on two test-pieces, including benzene dissociation from the protein lysozyme, where we calculate the dissociation pathway and timescales slower than milliseconds. Finally, by performing an analysis of residues contributing to the PIB, we predict the critical mutations in the system which would be most impactful on the stability of the crucial but ephemeral transition state. We believe this work marks a big step forward in the use of predictive artificial intelligence ideas for the sampling of biomolecules.

---

A key contributor to the rich and diverse functioning of molecular systems is the presence of myriad possible configurations. Instead of simply staying in the ground state, a given system can adopt one of many metastable configurations and stay trapped there for extended periods of time. It has been a longstanding dream to apply all-atom molecular dynamics (MD) simulations to learn what these metastable states are, their relative thermodynamic propensities, the pathways for moving between them, and associated kinetic constants. However there have been two central challenges in the use of MD for this purpose: (a) the large number of states and pathways for traversing between them, and (b) the inherent rare event nature of transition between states, wherein a simulation would simply be trapped in whichever metastable state it was started in. While multiple creative sampling methods^1^ and even ultra-specialized supercomputers^2^ have been introduced for tackling this timescale problem, the problem is not yet fully solved. For instance, a large class of sampling methods need an *a priori* sense of a reaction coordinate (RC), which is a low-dimensional summary of the many configurations and pathways.^3–7^ However this leads to an inherent coupled problem where one needs extensive sampling of rare events to learn the RC, but also needs to know the RC in the first place to perform sampling.

To address this problem, in this article our *ansatz* is that efficient sampling of energy landscapes of molecular systems has the same key underlying challenge as one faced by a fly as it goes about surviving^8^, or the human brain and retina trying to process how to catch a moving baseball^9^. Namely, given limited storage and computing resources, which memories to preserve and which ones to ignore in order to be best prepared for various possible future challenges? This can be paraphrased as the ability to rapidly learn a low-dimensional representation of a complex system that carries maximal information about its future state. Since storing and processing large amounts of information can be computationally and thus energetically expensive for the brain, it has been suggested and demonstrated that neurons in the brain separate predictive information from the non-predictive background in a way that by encoding and processing a minimum amount of relevant information, the brain can still be maximally prepared of future outcomes. The past–future (or Predictive) Information Bottleneck framework introduced and developed in many forms by different groups^8–11^ involves implementing such neuronal models from an information theoretic basis that can originally be traced back to Shannon’s rate distortion theory.

Here we interpret the reaction coordinate (RC) in molecular systems as such a past–future information bottleneck^10^. We develop a sampling method that for small biomolecules such as protein–ligand systems, simultaneously and with minimal use of human intuition, learns what this bottleneck is, its thermodynamics and its kinetics. The central idea is that not all features of the past carry predictive value for the future. A complex model can be made to be very predictive, however it will often obscure physical interpretability and also end up summarizing noise. In order to address this task, we set up an optimization problem and demonstrate how it can be solved through the use of the principle of variational inference^12^ implemented through deep neural networks. This makes it possible to learn a predictive information bottleneck^11^, which we interpret as the RC, that given a molecule’s past trajectory is maximally predictive of its future behavior. Our net product is an iterative framework on the lines of Ref.^13^ that starts from a short MD simulation, and given this data, makes an estimate of the RC, its Boltzmann probability, and its associated causal Green’s function valid for short times. This information is leveraged to perform systematically biased simulations with enhanced exploration of phase space, which can then be used to re-learn the RC along with its probability and propagator, and iterating between MD and variational inference until optimization is achieved. At this point we have converged estimates of the most informative degrees of freedom, associated metastable states and their equilibrium probabilities. Finally, through the use of a generalized transition state theory based framework on the lines of Ref.^14^, we recover the unbiased kinetics for moving between different metastable states along with their lifetimes.

We first demonstrate the method on sampling the metastable states in a small peptide. We then apply it to a problem of immense theoretical and practical relevance, by calculating the full dissociation process of benzene molecules from the L99A mutant of the 19 kDa protein T4 lysozyme^15,16^. In the last system, with use of all-atom MD simulations taking barely a few hundred nanoseconds in total, and with the minimal use of prior human intuition as in other related methods, we obtain accurate thermodynamic and kinetic information for a process that takes few hundred milliseconds in reality. Our simulations shed light on the complex interplay between protein flexibility and ligand movement, and predict, in good agreement with experiments, the residues whose mutations will have the strongest effect on the ligand dissociation mechanism. We believe our approach marks a big step forward in the use of fully-automated all-atom simulations for the study of complex molecular and biomolecular mechanisms.

## THEORY

### Principle of Past–Future Information Bottleneck

We formalize this problem in terms of a high-dimensional signal *χ*characterizing the state of a N-particle system under some generic set of thermodynamic conditions. We take *χ*as some *d* generalized coordinates or basis set elements, where 1 ≪ *d ≪ N.* Let the value of this signal measured at time *t*, or the “past”, be denoted by *X*_*t*_ and at time *t* + Δ*t*, or the “future” by *X*_*t*+Δ*t*_. We call Δ*t* the prediction time delay. We assume that *X*_*t*_ and *X*_*t*+Δ*t*_ are jointly distributed as per some probability distribution *P* (*X*_*t*_, *X*_*t*+Δ*t*_). The mutual information *I*(*X*_*t*_, *X*_*t*+Δ*t*_) (see Supplemental Information (SI) for this and other definitions) quantifies how much an observation at one instant of time *t* can tell us about an observation at another instant of time *t*+Δ*t*. Furthermore, in this article we restrict our attention to stationary systems, hence we omit the choice of time origin and write down *X*_*t*_ as *χ*and *X*_*t*+Δ*t*_ as *X*_Δ*t*_. The principle of Predictive information bottleneck (PIB)^10,11^ postulates a “bottleneck” variable *χ* which is related to *χ*by an encoder function *P* (*χ| X*). Given the bottleneck variable *χ*, predictions of the future *X*_Δ*t*_ can be made with a *decoder P* (*X*_Δ*t*_*|χ*). PIB says that the optimal bottleneck variable is one which is as simple as possible in terms of the past it needs to know, yet being as powerful as possible in terms of the future it can predict correctly. This intuitive principle can be formally stated through the optimization of an objective function ℒ which is a difference of two mutual informations:

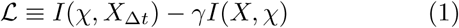

The above objective function quantifies the trade-off between complexity and prediction through a parameter *γ ∈* [0,1].

### Variational inference and neural network architecture

Typically, both the encoder *P* (*χ|X*) and the decoder *P* (*X*_Δ*t*_*|χ*) can be implemented by fitting deep neural networks^17^ to data in form of time-series of *X*. Our work stands out in three fundamental ways to typical implementations of the information bottleneck principle and in general of AI methods to sampling biomolecules^18–21^. Firstly, we use a stochastic deep neural network to implement the decoder *P* (*X*_Δ*t*_|*χ*), but use a simple deterministic linear encoder *P* (*χ|X*) (see Fig. 1). The simple encoder ensures that the information bottleneck or RC we learn is actually physically interpretable, which is notably hard to achieve in machine learning. On the other hand by introducing noise in the decoder, we can control the capacity of the model to ensure that the neural network can delineate useful feature from useless information instead of just memorizing the whole dataset. Secondly, now that our encoder is a simple linear model, we completely drop the complexity term in Eq. 1 and set *γ* = 0. Due to a reduced number of variables, this leads to a simpler and more stable optimization problem. Finally, the rare event nature of processes in biomolecules makes it less straightforward to use of information bottleneck/AI methods for enhanced sampling. Here we develop a framework on the lines of^13^ that makes it possible to maximize the objective function in Eq. 1 through the use of simulations that are progressively biased using importance sampling as an increasingly accurate information bottleneck variable is learnt.

**FIG. 1:**
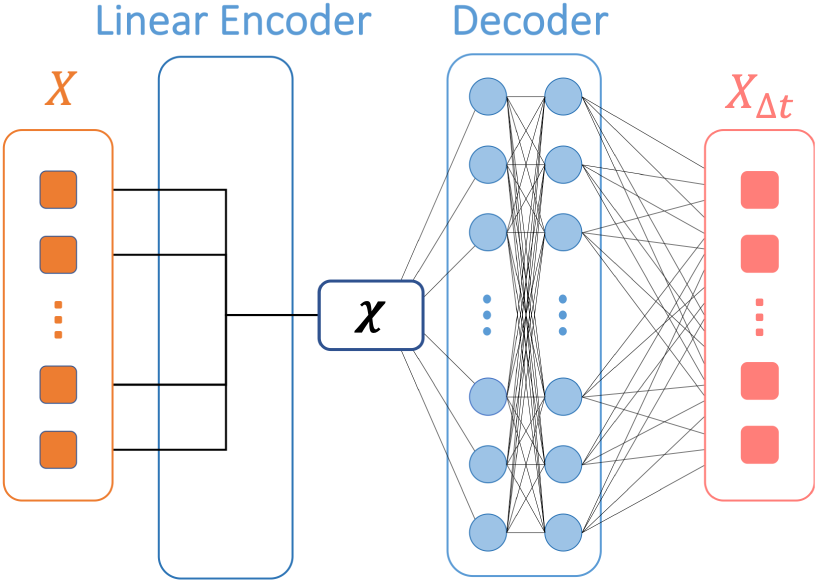
Network architecture using for learning predictive information bottleneck *χ*. The decoder *Q*(*X*_Δ*t*_ | *χ*) is a stochastic deep neural network, while the encoder *P* (*χ | X*) is of a simple deterministic and thus directly interpretable linear form.

Our typical starting point is an unbiased MD trajectory *χ*= {*X*^1^, *X*^2^,*…, X*^*M*^} with *M* data points. We want to develop a low-dimensional mapping *χ* of this high-dimensional space, that maximizes the objective function ℒ = *I*(*χ*(*X*), *X*_Δ*t*_). At the heart of this mutual information lies the calculation of the decoder *P* (*X*_Δ*t*_ *χ*), which can in principle be done *exactly* using Bayes’ theorem (SI). However this becomes impractical as soon as the dimensionality of *χ*increases, due to a fundamental problem common to statistical mechanics and machine learning: intractability of the partition function in high dimensions^22,23^.

Much of modern statistical physics and machine learning have attempted to surmount this problem with the advent of specialized methods^22,23^. The principle of variational inference is one such elegant and extremely powerful approach. Let’s consider a generic encoder given by some conditional probability *P*_*θ*_(*χ|X*) where *θ* is a set of parameters. Our objective then is to find the optimal RC or equivalently, the encoder *θ* which optimizes the PIB objective:

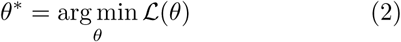

As mentioned above, this optimization problem is intractable for almost all cases of practical interest. However, it is possible to perform an approximate inference problem by assuming an approximate decoder *Q*_*Φ*_(*X*_Δ*t*_*|χ*) parametrized by the vector *Φ*. For any choice of *Φ*, we make a straightforward use of Gibbs’ inequality^12^ to write down (SI):

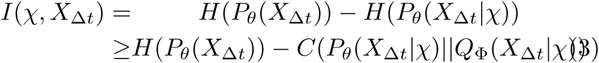

Here H and C denote Shannon entropy and cross entropy respectively. Take note that the first term in Eq. 3 is independent of our model parameters and hence can be completely ignored from the optimization. Focusing on the second term in Eq. 3, we thus obtain a variational lower bound on the predictive information bottleneck objective function:

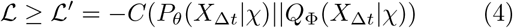

Thus ℒ *′* is a tractable lower bound bound to the true Predictive Information Bottleneck objective function ℒ, that involves a variational approximation through the trial decoder parametrized by Φ. It has a simple physical interpretation. We are attempting to learn a decoder probability function *Q* that mirrors the actual Bayesian inverse probability function *P* in terms of predicting the future state *X*_Δ*t*_ of the system, given knowledge of the RC *χ*. The difference between the two probability distributions is calculated as a cross-entropy. By maximizing the right hand side of Eq. 4 simultaneously with respect to the decoder and encoder parameters Φ and *θ* respectively, we can then solve the actual optimization problem posited in Eq. 4 rigorously and identify the optimal RC.

It is clear that a model of a dynamical system *χ*that attempts to capture just its stationary probability *P* (*X*) will be less informative and useful than one that captures the joint past-future probability distribution *P* (*X, X*_Δ*t*_). This is simply because the stationary probability can always be calculated by integrating *P* (*X, X*_Δ*t*_) over future outcomes *X*_Δ*t*_. What is however less clear is the choice of the time-delay Δ*t*^24^. In biomolecular systems, it is likely that there will be a hierarchy of time-scales and thus time-delays relevant to different types of structural and functional details. In principle, our formulation allows us to probe these various time-delays in a systematic manner. Here, for the purpose of enhanced sampling, we propose an approach for selecting Δ*t* that is rooted in the reactive flux formalism of chemical kinetics^25–27^. This formalism applies to any system with stochastic transitions on a network of microstates with arbitrary, complex connectivity. Summarily, it states that the correlation function for a trajectory’s population in any given state can be partitioned into 3 parts: (a) an initial transient, inertial part, (b) an eventual exponential decay, and (c) an intermediate plateau region between the very short-time and long-time parts. A key insight from this formalism is that capturing (c), i.e. the plateau parts of a system’s state to state dynamics accurately is necessary and sufficient to capture the temporal evolution at any timescale. By paraphrasing this argument in the context of the present work, we propose to learn our PIB model for gradually increasing values of the predictive time-delay Δ*t*, and stop when the calculated bottleneck variable coverges (see numerical examples in **Results**), identifying it as the plateau from reactive flux formalism.

### Variational inference on unbiased and biased trajectories

We now show how to calculate *ℒ′*in practice. For a given unbiased trajectory {*X*^1^, *X*^2^,*…, X*^*M*+*k*^}with large enough *M*, we can easily show (SI):

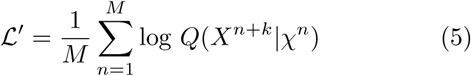

where *χ*^*n*^ is sampled from *P* (*χ | X*^*n*^) and the time interval between *X*^*n*^ and *X*^*n*+*k*^ is Δ*t*. For practical rare event systems however, a typical MD trajectory will be trapped in the state where it was started. Here we use our current best estimate of the PIB to perform importance sampling of the landscape, so that the system is more likely to sample different regions in configuration space, and use this enhanced sampling to iteratively improve the quality of the RC. However, the data so generated is biased per definition, and we need to reweight out the effect of the bias. We suppose that along with the time series {*X*^1^, *X*^2^,*…, X*^*M*+*k*^} we also have been provided the corresponding time-series for the bias *V* applied to the system {*V* ^1^, *V* ^2^,*…, V* ^*M*+*k*^}. We can then use the principle of importance sampling^28^ to write our PIB objective function *ℒ′*as follows (SI):

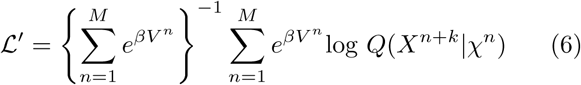

where *β* is the inverse temperature. The above equation is however approximate, as it makes the assumption *P*_*biased*_(*X*^*n*+*k*^ | *χ*^*n*^) ≈ *P*_*unbiased*_(*X*^*n*+*k*^ | *χ*^*n*^). This is exact as Δ*t* 0, and can be expected to be reasonably valid for small enough Δ*t*. Namely, for small Δ*t*, we expect that the system in time interval Δ*t* on an average would not have diffused too far from its starting position at the beginning of that interval. If the bias varies smoothly enough that its natural variation length scale is smaller or comparable to this diffusion distance, then for small enough Δ*t* we can indeed make the aforementioned approximation. This means that we select the smallest possible Δ*t* at which the RC estimate plateaus. In principle a more exact version of Eq. 6, one which exactly corrects the effect of bias even on the short time propagator of the system, can be derived and used however the computational costs for such procedures can be formidable enough to make them impractical for our purpose.

### Patching it all

We now state our complete enhanced sampling algorithm, that accomplishes in a seamless manner, the identification of the RC together with the sampling of its thermodynamics and kinetics, generalizing the framework from Ref.^13^. The first step is to perform an initial round of unbiased MD simulation. This trajectory, expressed in terms of *d* order parameters or basis functions {*s*_1_,*…, s*_*d*_} (where 1 ≪*d ≤ N*), is fed to a deep learning module (Fig. 1). The deep learning module implements the optimization of *ℝ′*in Eq. 6 through the use of multi-layer feed-forward neural network for the stochastic decoder *Q*, and a physically interpretable linear map for the deterministic encoder *P* (Fig. 1). Unlike the decoder, the encoder has no noise term and always maps {*s*_1_, *s*_2_,*…, s*_*d*_} to ∑_*i*_ *c*_*i*_*s*_*i*_, where {*c*_*i*_} denote the weights of different order parameters. We perform this optimization for gradually increasing values of the predictive time-delay Δ*t*, and estimate RC *χ* (given by the values of the weights *c*_*i*_) as seen by the first “plateau” in terms of when it ceases to depend on choice of Δ*t*. This value of Δ*t* is then kept constant for different rounds of our protocol. At this point we have an initial estimate of *χ* and also its unbiased probability distribution *P*^*u*^(*χ*). These are both used to construct a bias potential *V*_*bias*_(*χ*) for the next iteration of MD:

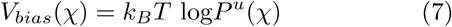

where *k*_*B*_ is Boltzmann’s constant and *T* is the temperature. With this bias potential added to the original Hamiltonian of the system, we run a biased MD simulation. This explores an increased amount of configuration space since we have applied a bias along our estimated slow degree of freedom, viz. the PIB or the RC. This next round of MD trajectory is once again fed to the deep learning module, but this time each data point carries a weight 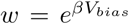 to compensate for the applied bias. This now identifies an improved RC *χ* and its unbiased probability through the use of importance sampling:

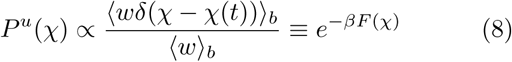

where the subscript *b* denotes sampling under a biased ensemble with weight 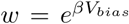 and *F* (*χ*) is the free energy along *χ*. From here, using the bias as –*F* (*χ*) our algorithm can now enter into further iterations of –MD–deep learning–MD–… This looping continues until both the RC *χ* and the free energy estimate *F* (*χ*) along the RC have converged. We have thus obtained an optimized reaction coordinate and its Boltzmann probability density, or equivalently the free energy. Through these we can directly demarcate the relevant metastable states and quantify their relative propensities. Furthermore, we can also calculate the transition rates for moving between these metastable states. The central idea is to keep all transition states between the different metastable states, as identified through the RC, devoid of any bias. As we show in examples, this can be easily achieved when implementing Eq. 8, by ensuring that any barriers in the unbiased probability distribution of the estimated RC are completely bias-free. Once we have done this, we take into account that by virtue of it being the PIB, the RC already encapsulates any relevant, predictive modes in the system. Thus the hidden modes and barriers which have invariably been corrupted through the addition of such a bias do not have any predictive power for the dynamics of the system, and are thus not relevant to the process at hand. This then implies that (i) the biased dynamics preserves the state-to-state sequence one would have seen with unbiased dynamics, and (ii) through the use of a simple time rescaling calculation (see SI and Ref.^14^) we can the calculate the acceleration of rates achieved through biased simulations. Finally, we can perform self-consistency checks for the reliability of the rescaled kinetics by analyzing the unbiased lifetimes for robustness with precise choice of biasing protocols (SI).

## RESULTS

We now demonstrate the use of the PIB framework with two biomolecular case-studies, in both of which we simultaneously learn the RC, the free energy and kinetic rate constants. In each case the RC *χ* is constructed as a linear combination *χ* = ∑_*i*_ *c*_*i*_*s*_*i*_, where {*c*_*i*_} denote the weights of different pre-selected order parameters *{s*_1_, *s*_2_,*…, s*_*d*_*}*.

### Conformation transitions in a model peptide

First we consider the well-studied alanine dipeptide system (Fig. 2(a)). This system, as characterized by its Ramachandran dihedral angles, can exist in different metastable states with varying stabilities and hard-to-cross intermediate barriers. However, due to its small size (22 atoms) it serves as a reliable benchmark where we can perform longer than microsecond unbiased MD simulations to compare our PIB calculations against.

**FIG. 2:**
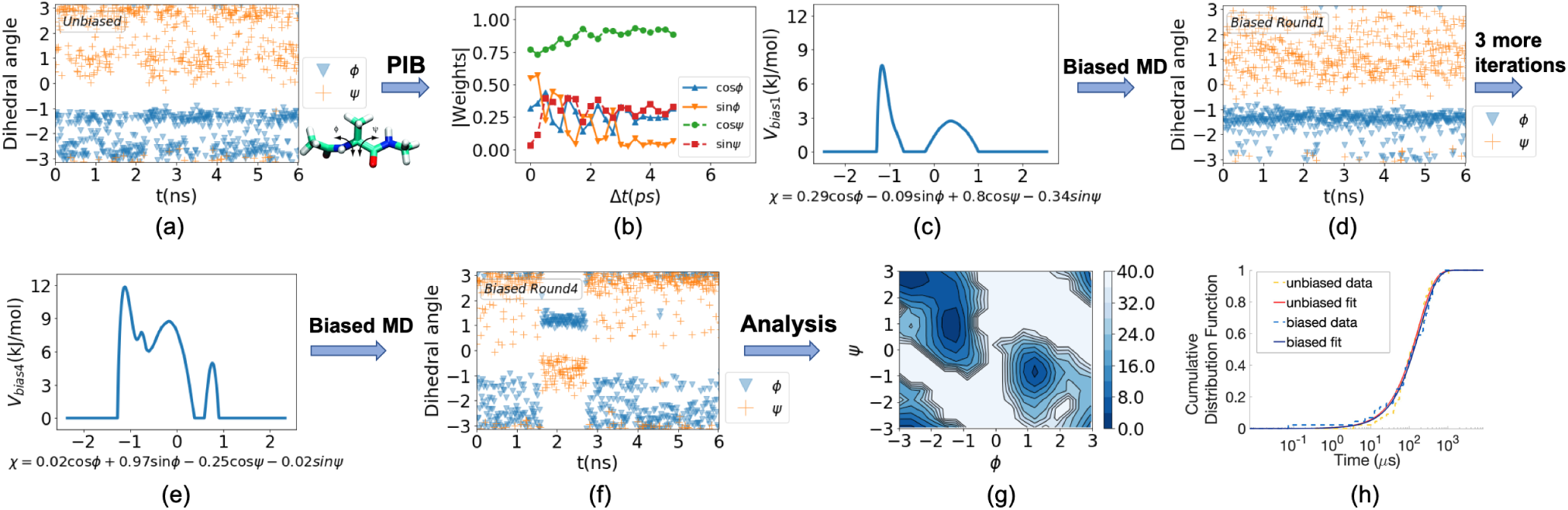
(a) Unbiased simulation trajectory. The alanine dipeptide molecule is shown in inset. (b) Absolute weights for different order parameters in the first training round as a function of the predictive time delay Δt. (c-f) Free energy along the adaptively learnt RC along with the corresponding biased trajectories for different training rounds. (g) Free energy along *ϕ, ψ* after the RC has converged. (h) Kinetics from the post-training biased runs as well as reference unbiased runs. The two are essentially indistinguishable.

Here we choose {cos *ϕ*, sin *ϕ*, cos *ψ*, sin *ψ*} as our order parameters, where *φ* and *ψ* are the backbone dihedral angles (marked in the inset of Fig. 2(a)). By taking trigonometric functions of dihedral angles, we avoid problems related to periodic boundary conditions. The PIB protocol used here is shown in Fig. 2. In the initial round, we perform a short unbiased MD simulation (see Fig. 2(a) for trajectory and SI for technical details). The neural network decoder consists of two hidden layers, with 128 neurons in each layer. To safeguard against spurious local minima solution learnt by the neural network, we perform independent training runs with random initial weights for order parameters. The RC is then determined as the linear encoder of the trained neural network with smallest loss function during training. In Fig. 2(b), we show how the weights rapidly converge as functions of predictive time delay Δ*t* and reach a plateau in less than 2 ps. We then set Δ*t* = 2 ps in all future rounds. Fig. 2(c) shows the RC *χ* as well as the bias *V* (*χ*) learnt along it to be used in the next round of MD. With this biasing potential, we perform biased MD simulation as shown in Fig. 2(d). This trajectory through the use of Eq. 8 leads to a more complicated bias structure as shown in Fig. 2(e) along with the improved RC *χ*. Biased simulation with this new RC and bias as shown in Fig. 2(f) finally leads to escape from the starting metastable state. The final obtained RC is: *χ* = 0.02 cos *ϕ* + 0.97 sin *ϕ –* 0.25 cos *ψ –* 0.02 sin *ψ*. It is known for alanine dipeptide that *ϕ* is more relevant than *ψ* for capturing the conformational transitions, and our PIB based RC estimate agrees with that. The shift in weights of order parameters across different rounds (SI) reflects how our iterative scheme finds the optimal RC: a better RC provides better sampling and using the information from the better sampling we can learn a new RC which can further enhance the sampling, and so on.

Now that we have achieved back-and-forth motion in terms of the rare event we intended to study, we use this final RC and bias to perform multiple sets of longer simulations with no further refinement of the RC. This yields the free energy surface (defined as –*k*_*B*_*T* log*P* (*ϕ, ψ*) where *P* is the unbiased Boltzmann probability) as shown in Fig. 2(g). This is in excellent agreement with previously published benchmarks for this system^28^. At the same time, we use the acceleration factor to rescale the biased time back to the unbiased time. In Fig. 2(h) we show the cumulative distribution functions of the first passage time from the deeper basin as obtained through this approach, and through much longer unbiased MD runs which are feasible given the small size of this system. The distribution functions and their best-fit Poisson curves are nearly indistinguishable, and lead to excellent agreement in values of the escape rate constant from the deeper basin, given by *k* = 5.2 *±* 0.8*µs*^-1^ and 5.8*±*0.9*µs*^-1^ respectively for biased and unbiased simulation respectively.

### Benzene dissociation from T4-L99A lysozyme

We now apply our framework to a very challenging and important test case, namely the pathway and kinetic rate constant of benzene dissociation from the protein T4-L99A lysozyme in all-atom resolution^15,16^. We also demonstrate how the RC calculated through our approach can be directly used to perform a sensitivity analysis of the protein, and predict the most important residues whose mutations could have a significant affect on the stability of the protein-ligand complex. Such an analysis has direct relevance to predicting, for instance, the mutations in a protein which could lead to a pharmacological drug losing its efficacy. For this problem we choose 11 fairly arbitrary order parameters denoted {*s*_1_,*…, s*_11_}. Eight of these are ligand-protein distances while three are intra-protein distances (see SI for order parameter details). The RC is learnt as a linear combination of these order parameters, namely *χ* = ∑_*i*_ *c*_*i*_*s*_*i*_.

For this problem as well we start with a short unbiased MD simulation. As shown in Fig. 3, the weights of different order parameters in the RC learnt from this trajectory change as a function of the predictive time-delay Δ*t*, but converge quickly. On the basis of this plot, we set Δ*t* = 2*ps* for all further calculations. We then iterate – using the same neural network architecture as for alanine dipeptide (Fig. 1) – between rounds of learning an iteratively improved RC *χ*_1_ together with its probability distribution, and running biased MD using the iterations RC and probability distribution as bias *V*_1_(*χ*_1_) (Eq.7). After 9 rounds, we find that the bias saturates as a function of training rounds. That is, no further enhancement in egodicity is achieved by performing additional rounds of the aforementioned iteration. This corresponds to the system reaching configuration where the previous PIB ceases to be effective. To learn a new PIB, we use the “washing out” trick from^29^ to learn a second RC *χ*_2_ conditioned on our knowledge of the first RC *χ*_1_. In the next few rounds of learning–MD iterations, we (a) keep *χ*_1_ and *V*_1_(*χ*_1_) fixed, and (b) do not account for *V*_1_(*χ*_1_) when using Eq.8. Through this we learn a bias *V* (*χ*_1_, *χ*_2_) = *V*_1_(*χ*_1_)+ *V*_2_(*χ*_2_). In principle we can lift this assumption and learn more complicated non-separable *V* (*χ*_1_, *χ*_2_). In a few rounds of training *χ*_2_, we observed spontaneous disassociation of the ligand from the protein. We are now ready to use the RC (*χ*_1_, *χ*_2_) (shown in Fig. 4) and its bias *V* (*χ*_1_, *χ*_2_) learnt to directly study the pathway and kinetics of ligand dissociation.

**FIG. 3:**
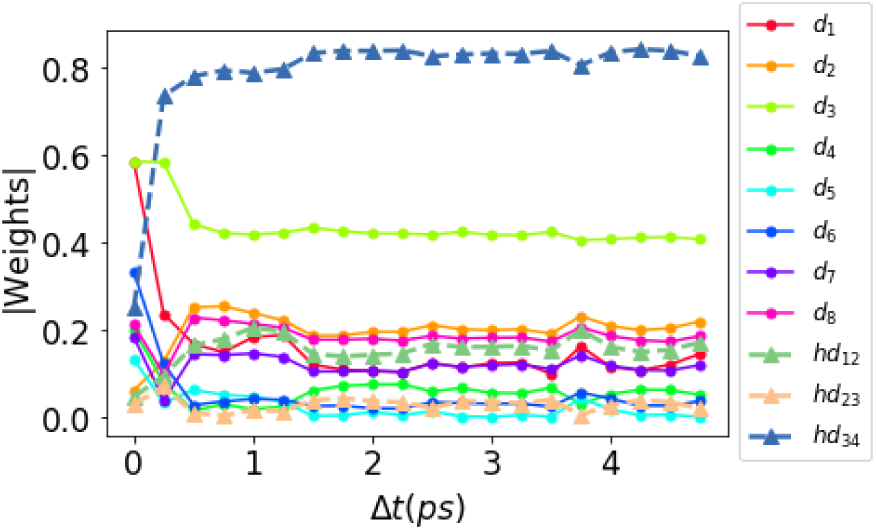
Benzene-lysozyme dissociation: The absolute value of weights for 11 order parameters in the first round as a function of predictive time delayously

**FIG. 4:**
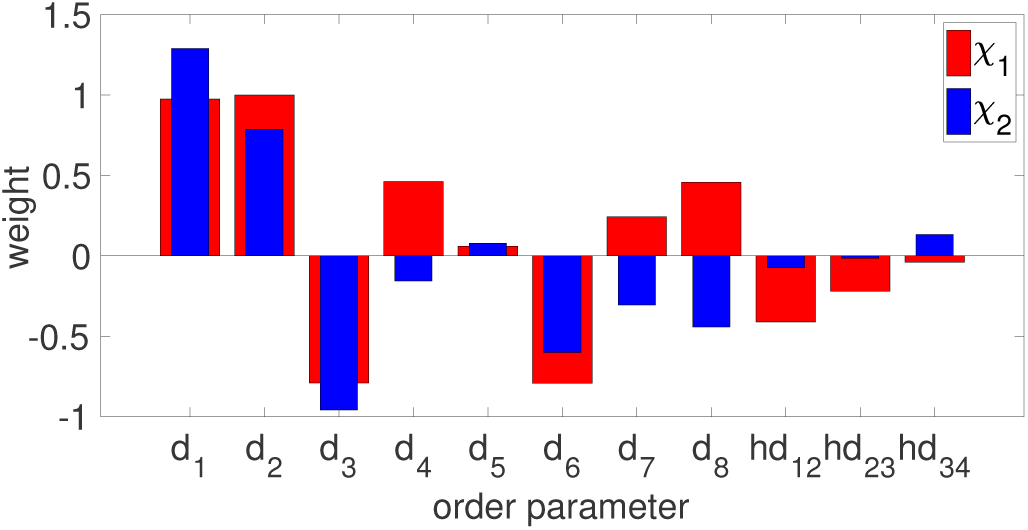
The 2-component Predictive Information Bottleneck for benzene-lysozyme dissociation, where the optimized weights for different order parameters are illustrated, after scaling all weights to keep *c*_1_ = 1 in *χ*_1_.

For this we launch 20 independent biased simulations using (*χ*_1_, *χ*_2_) as RC and *V*_1_(*χ*_1_) + *V*_2_(*χ*_2_) as bias. By calculating the acceleration factor, we can recover the original timescale of the first passage time. As we show in SI, we fit the cumulative distribution function to a Poisson process and get an escape rate constant of 3.3 *±* 0.8*/s*, which is in good agreement with other simulation methods^30–32^. We also obtain a range of free energies viewed as functions of different order parameters (SI). These are in excellent agreement with previously published results, especially the ones using same forcefields^29–32^.

### Predicting critical residues

On the basis of the predictive information bottleneck that we have now calculated, we can directly predict which protein residues have the most critical effect on the benzene-lysozyme transition state. To do so, our guiding principle is that the residues which carry higher mutual information with the predictive information bottleneck are more likely to have an impact on the stability of the system, for instance, if these residues were to be mutated. By performing a scan of the mutual information between the predictive information bottleneck and the backbone dihedral angles of different residues, we can rank them as being most critical to least critical (see SI for further details of the calculation set-up). As shown in Fig. 5, some of the important residues are (in order of decreasing relevance): Ser136, Lys135, Asn132, Leu133, Ala134, Phe114, Val57, Asp20, Leu118 and Val131. These residues can be classified in three broad groups: (a) residue 114, and residues 131–136: together these contribute to breathing movement between the two helices through which the ligand leaves, (b) residue 118, which is the major hydrophobic interaction in the bound state, and (c) residues 20 and 57, which lie in different disordered regions between ordered parts of the protein, and have no obvious interpretation. The roles of groups (a) and (b) have been hinted at in previous works^32–34^ and are thus yet another validation of our approach. It remains to be seen if group (c) indeed has biophysical relevance possibly through a long-range allosteric communication pathway, or is just noise from our calculations.

**FIG. 5:**
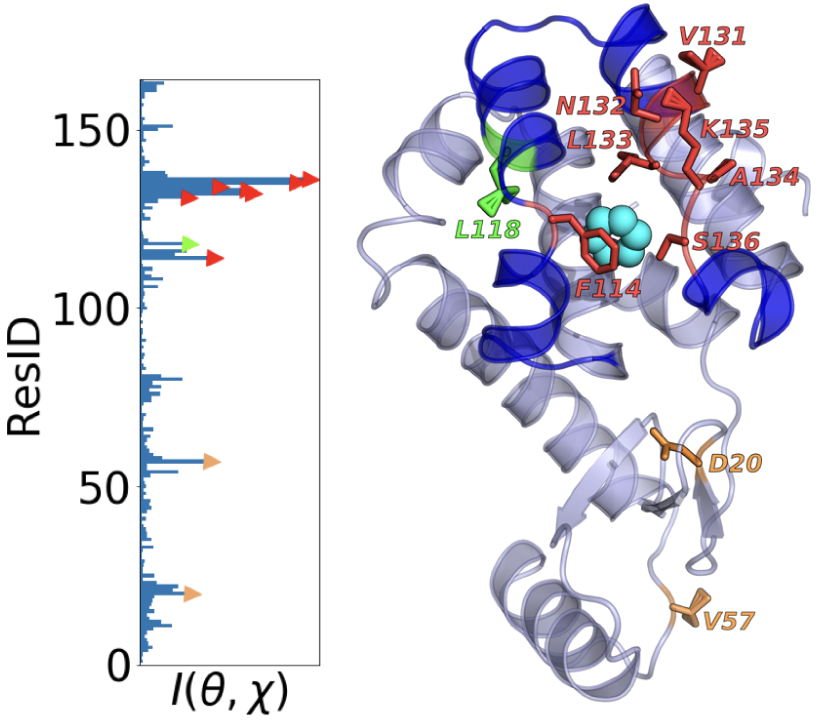
Critical residue analysis for benzene-lysozyme complex. The plot on left shows for every residue the maximal mutual information between the PIB and either of the Ramachandran angles *ϕ, ϕ* of that residue. The top 10 residues are highlighted through markers and in the right plot, illustrated relative to the ligand in a typical intermediate pose.

## DISCUSSION

In this work we have introduced a new framework for the simultaneous sampling of the reaction coordinate, free energy and rate constants in biomolecules with rare events. Our work is grounded in the Predictive Information Bottleneck framework, which is an information theoretic approach for building minimally complex yet maximally predictive models from data. Such a framework has previously been found useful for modeling fruit fly movement and human vision. Here we exploit the commonality between these diverse problems and that of sampling complex biomolecular systems, namely the need to quickly predict the future state of a system given noisy and high-dimensional information. Our method implements this framework through the use of a unique “linear encoder–stochastic decoder” model, where the latter is a deep neural network with inbuilt noise. Here we demonstrated the applicability of the method by studying conformational transitions in a model peptide and ligand dissociation from a protein, in explicit water and all-atom resolution. Through extremely short and computationally cheap simulations, we obtained thermodynamic and kinetic observables for slow biomolecular processes in excellent agreement with other methods, experiments and long unbiased MD. Last but not the least, by virtue of having captured the most predictive degrees of freedom in the system, we could also make, arguably for the firs time, direct predictions of how protein sequence can impact dissociation dynamics - namely, which mutations in the protein would be most deleterious to the dissociation process. We believe this work marks an important step ahead in computer simulation of molecules, and should be useful to different communities for robust, reliable studies of rare events.

## ACKNOWLEDGMENTS

We thank Deepthought2, MARCC and XSEDE (projects CHE180007P and CHE180027P) for computational resources used in this work. PT would like to thank the University of Maryland Graduate School for financial support through the Research and Scholarship Award (RASA).

## References

1. P. Tiwary and A. van de Walle, in Multiscale Materials Modeling for Nanomechanics (Springer, 2016) pp. 195–221.

2. K. Lindorff-Larsen, S. Piana, R. O. Dror, and D. E. Shaw, Science 334, 517 (2011).

3. A. Ma and A. R. Dinner, J. Phys. Chem. B 109, 6769 (2005).

4. R. B. Best and G. Hummer, Proc. Natl. Acad. Sci. 102, 6732 (2005).

5. A. Berezhkovskii and A. Szabo, The Journal of chemical physics 122, 014503 (2005).

6. P. Tiwary and B. J. Berne, Proc. Natl. Acad. Sci. 113, 2839 (2016).

7. P. Tiwary and B. J. Berne, J. Chem. Phys. 145, 054113 (2016).

8. G. J. Berman, W. Bialek, and J. W. Shaevitz, Proceedings of the National Academy of Sciences 113, 11943 (2016).

9. S. E. Palmer, O. Marre, M. J. Berry, and W. Bialek, Proc. Natl. Acad. Sci. 112, 6908 (2015).

10. N. Tishby, F. C. Pereira, and W. Bialek, arXiv preprint physics/0004057 (2000).

11. S. Still, Entropy 16, 968 (2014).

12. D. J. MacKay and D. J. Mac Kay, Information theory, inference and learning algorithms (Cambridge university press, 2003).

13. J. M. L. Ribeiro, P. Bravo, Y. Wang, and P. Tiwary, J. Chem. Phys. 149, 072301 (2018).

14. P. Tiwary and M. Parrinello, Phys. Rev. Lett. 111, 230602 (2013).

15. A. E. Eriksson, W. A. Baase, X.-J. Zhang, D. W. Heinz, M. Blaber, E. P. Baldwin, and B. W. Matthews, Science 255, 178 (1992).

16. V. A. Feher, E. P. Baldwin, and F. W. Dahlquist, Nature Structural and Molecular >Biology 3, 516 (1996).

17. A. A. Alemi, I. Fischer, J. V. Dillon, and K. Murphy, arXiv preprint arXiv:1612.00410 (2016).

18. A. Mardt, L. Pasquali, H. Wu, and F. Noé, Nat. Comm. 9, 5 (2018).

19. W. Chen and A. L. Ferguson, Journal of computational chemistry 39, 2079 (2018).

20. C. Wehmeyer and F. Noé, The Journal of Chemical Physics 148, 241703 (2018).

21. M. M. Sultan, H. K. Wayment-Steele, and V. S. Pande, arXiv preprint arXiv:1801.00636 (2018).

22. I. Goodfellow, Y. Bengio, A. Courville, and Y. Bengio, Deep learning, Vol. 1 (MIT press Cambridge, 2016).

23. N. Goldenfeld, Lectures on phase transitions and the renormalization group (CRC Press, 2018).

24. C. Wehmeyer and F. Noé, arXiv preprint arXiv:1710.11239 (2017).

25. B. J. Berne, M. Borkovec, and J. E. Straub, J. Phys. Chem. 92, 3711 (1988).

26. J. A. Montgomery Jr, D. Chandler, and B. J. Berne, The Journal of Chemical Physics 70, 4056 (1979).

27. C. Dellago, P. G. Bolhuis, and D. Chandler, The Journal of chemical physics 110, 6617 (1999).

28. O. Valsson, P. Tiwary, and M. Parrinello, Ann. Rev. Phys. Chem. 67, 159 (2016).

29. J. M. L. Ribeiro and P. Tiwary, Journal of chemical theory and computation (2019), 10.1021/acs.jctc.8b00869.

30. Y. Wang, J. M. Martins, and K. Lindorff-Larsen, Chemical science 8, 6466 (2017).

31. Z. Smith, D. Pramanik, S.-T. Tsai, and P. Tiwary, J. Chem. Phys. 10.1063/1.5064856.

32. J. Mondal, N. Ahalawat, S. Pandit, L. E. Kay, and P. Vallurupalli, PLoS computational biology 14, e1006180 (2018).

33. G. Bouvignies, P. Vallurupalli, D. F. Hansen, B. E. Correia, O. Lange, A. Bah, R. M. Vernon, F. W. Dahlquist, D. Baker, and L. E. Kay, Nature 477, 111 (2011).

34. M. D. Collins, G. Hummer, M. L. Quillin, B. W. Matthews, and S. M. Gruner, Proceedings of the National Academy of Sciences 102, 16668 (2005).

