## Supplement for "Past–future information bottleneck framework for simultaneously sampling biomolecular reaction coordinate, thermodynamics and kinetics"

#### This PDF file includes:

Supplementary text
Figs. S1 to S5
Table S1
References for SI reference citations

### Supporting Information Text

### Definitions

Here we define various terms introduced in the main text as well. In all of these we use $P(X)$, $P(X,Y)$ respectively to denote the probability distribution of a random variable $X$ and the joint probability distribution of two random variables $X$ and $Y$. Other variables are the same as defined in the main text.

1. **Mutual information**

   This is a commonly used information theoretic measure to describe how much information is shared between two random variables $X$ and $Y$. It is defined as:

$$I(X,Y) = \int P(X,Y) \ln \frac{P(X,Y)}{P(X)P(Y)} dX dY \quad [1]$$

2. **Shannon entropy**

   The Shannon entropy for a random variable $X$ is defined as:

$$H(X) = -\int P(X) \ln P(X) dX \quad [2]$$

3. **Cross entropy**

   The cross entropy between two probability distributions $P(X)$ and $Q(X)$ is given by:

$$C(X,Y) = -\int P(X) \ln Q(X) dX \quad [3]$$

4. **Kullback-Leibler Divergence**

   The Kullback-Leibler (KL) $D_{KL}(P||Q)$ divergence between two probability distributions $P(X)$ and $Q(X)$ is given by:

$$D_{KL}(P||Q) = \int P(X) \ln \frac{P(X)}{Q(X)} dX \quad [4]$$

5. **Exact decoder definition**

   The exact decoder can be defined as the the conditional probability distribution of $X_{\Delta t}$ given a RC $\chi$. It can be calculated by using Bayes theorem:

$$P(X_{\Delta t}|\chi) = \frac{P(\chi, X_{\Delta t})}{P(\chi)} = \frac{\int P(\chi|X)P(X, X_{\Delta t})dX}{\int\int P(\chi|X)P(X, X'_{\Delta t})dX dX'_{\Delta t}} \quad [5]$$

6. **Acceleration factor**

   As shown in (1), and under the conditions detailed there, it is possible recover the unbiased timescales from biased simulations through the calculation of a simple acceleration factor. If the biased simulation time is $t$, the associated unbiased timescale $\tau$ can be calculated by:

$$\tau = \int_0^t e^{\beta V[\chi(t')]} dt' = \sum_{n=1}^{M} e^{\beta V^n} (t_n - t_{n-1}) \quad [6]$$

   where $V(\chi(t'))$ is the bias experienced by the system at time $t'$, constructed as a function of the RC $\chi$. This can equivalently be calculated as a discrete sum over the $M$ integration time-steps.

### Derivations

Here we provide the missing details of various derivations given in the main text.

1. **Gibbs's inequality and variational lower bound**

   The bottleneck function $\mathcal{L}$ defined in the main text for our neural network architecture is given by a difference of two Shannon entropies(2):

$$\mathcal{L} = I(\chi, X_{\Delta t}) = H(P(X_{\Delta t})) - H(P_\theta(X_{\Delta t}|\chi)) \quad [7]$$

Gibbs's inequality(2) guarantees that that the KL-divergence between two probability distributions is always larger than 0. We thus have:

$$
\begin{aligned}
D_{KL}(P_\theta(X_{\Delta t}|\chi))||Q_\Phi(X_{\Delta t}|\chi)) &\equiv \int\int P_\theta(X_{\Delta t},\chi)\ln\frac{P_\theta(X_{\Delta t}|\chi)}{Q_\Phi(X_{\Delta t}|\chi)}dX_{\Delta t}d\chi \\
&= \int\int P_\theta(X_{\Delta t},\chi)\ln P_\theta(X_{\Delta t}|\chi)dX_{\Delta t}d\chi - \int\int P_\theta(X_{\Delta t},\chi)\ln Q_\Phi(X_{\Delta t}|\chi)dX_{\Delta t}d\chi \\
&= -H(P_\theta(X_{\Delta t}|\chi)) + C(P_\theta(X_{\Delta t}|\chi), Q_\Phi(X_{\Delta t}|\chi)) \geqslant 0
\end{aligned} \quad [8]
$$

Only in the limit that our approximate decoder is exactly the same as the exact inverse-Bayes decoder, $D_{KL}(P_\theta||Q_\Phi)$ equals 0. By combining Eq.7 and Eq.8 , we get the relationship:

$$
\mathcal{L} \geqslant \mathcal{L} - D_{KL}(P_\theta(X_{\Delta t}|\chi))||Q_\Phi(X_{\Delta t}|\chi)) = H(P(X_{\Delta t})) - C(P_\theta(X_{\Delta t}|\chi), Q_\Phi(X_{\Delta t}|\chi)) \equiv H(P(X_{\Delta t})) + \mathcal{L}' \quad [9]
$$

Here, $H(P(X_{\Delta t}))$ only depends on the data set and is independent of the parametrization of the encoder and the decoder. Thus the term $H(P(X_{\Delta t})$ can be completely ignored while optimizing the parameters $\theta$ and $\Phi$. Maximizing the objective function $\mathcal{L}' = -C(P_\theta(X_{\Delta t}|\chi))$ is then equivalent to maximizing the lower bound of $\mathcal{L}$, which is the expression stated in Eq.4 in the main text.

2. **PIB objective $\mathcal{L}'$ for unbiased trajectory**

For a unbiased trajectory $\{X^1, X^2, ..., X^{M+k}\}$, for given $\theta$, we can get corresponding $\{\chi^1, \chi^2, ..., \chi^M\}$ using $\chi_i = \sum_i c_i s_i$. We also have a sampling of the states of the system after corresponding $\Delta t$ intervals: $\{X^{1+k}, X^{2+k}, ..., X^{M+k}\}$. Together the pair $(\chi_i, X_{i+k})$ sampled from the dataset follows the distribution $P_\theta(X_{\Delta t}|\chi)$. With this, we have:

$$
\mathcal{L}' = -C(P_\theta(X_{\Delta t}|\chi), Q_\Phi(X_{\Delta t}|\chi)) = \int\int P_\theta(X_{\Delta t},\chi)\ln Q_\Phi(X_{\Delta t}|\chi)dX_{\Delta t}d\chi = \frac{1}{M}\sum_{n=1}^{M}\log Q(X^{n+1}|\chi^n) \quad [10]
$$

3. **PIB objective $\mathcal{L}'$ for biased trajectory**

For a biased trajectory $\{X^1, X^2, ..., X^M\}$ with corresponding biasing potential values $\{V^1, V^2, ..., V^M\}$, the unbiased probability distribution of $X$ can be calculated by:

$$
P(X) = \frac{\sum_i \delta(X^i - X)e^{\beta V^i}}{\sum_i e^{\beta V^i}} \quad [11]
$$

The encoder $P(\chi|X)$ and decoder $P(X_{\Delta t}|\chi)$ are taken to be independent of the bias. The first assumption is strictly true, while the second is valid for small enough $\Delta t$ as explained in the main text. Therefore

$$
\mathcal{L}' = \int\int P_\theta(X_{\Delta t}|\chi)P(\chi)\ln Q_\Phi(X_{\Delta t}|\chi)dX_{\Delta t}d\chi = \left\{\sum_{n=1}^{M} e^{\beta V^n}\right\}^{-1}\sum_{n=1}^{M} e^{\beta V^n}\log Q(X^{n+1}|\chi^n) \quad [12]
$$

### Mutual information calculation for critical residue prediction

We use the backbone dihedral angles to describe the motion of each residue. Here we denote them as $\phi_i, \psi_i$, where $i$ is the index of a particular residue. We use $I(\theta,\chi)$ to quantify the correlation between dihedral angle $\theta$ (where $\theta$ could be $\phi$ or $\psi$) and the unbinding process, where $I(\theta,\chi)$ is mutual information between a dihedral angle $\theta$ and the RC $\chi$.

For each residue, we have two dihedral angles. At the same time, for one dihedral angle, we can calculate its mutual information with $\chi_1$ or $\chi_2$. So we have four quantities for each residue( $I(\phi,\chi_1)$,$I(\psi,\chi_1)$, $I(\phi,\chi_2)$ and $I(\psi,\chi_2)$ ). We rank the importance of each residue by the maximum of the four quantities. Here we only consider the parts of trajectory that is bias-free($\chi_1 > 0.4$ and $\chi_2 > 0.3$). Because we require that the energy barriers between metastable states should be bias-free to ensure we can get the correct reweighted dynamics of the unbinding process. However, we only guarantee that the main energy barrier between bound and unbound state has zero bias when we perform biased MD simulation while bias can still be added on barriers between other metastable states. Looking at the unbiased region all us to reduce the influence of the biasing potential on the dynamics of the system and focus more on the transition states.

### Simulation set-up for MD

**A. Alanine dipeptide in vacuum .** We follow (3) to set up our simulation for alanine dipeptide in vacuum. The simulations are performed with the software GROMACS 2016/GROMACS 5.0, patched with PLUMED 2.4. We constrain bonds involving hydrogens using the LINCS algorithm and employ an integration time-step of 0.002 ps. The temperature is kept constant at 300K using the velocity rescaling thermostat (relaxation time of 0.1 fs). We employ no periodic boundary conditions and no cut-offs for the electrostatic and non-bonded Van der Waals interactions.

**B. L99A T4L-benzene.** We follow (4) to set up our simulation for L99A T4L-benzene. The simulations are performed with the software GROMACS 2016/GROMACS 5.0 patched with PLUMED 2.4. The simulations are done with the constant number, pressure, temperature (NPT) ensemble with temperature 298 K and pressure 1.0 bar. Constant pressure is maintained using Parrinello-Rahaman barostat while the constant temperature is maintained using the v-rescale thermostat (modified Berendsen thermostat). The simulation box with periodic boundary condition is filled with TIP3P water. The side lengths of the box are 10Å and there are around $10,000$ water molecules. The interaction is described by the force field CHARMM22*. The integration time step here as well was taken to be 2 fs.

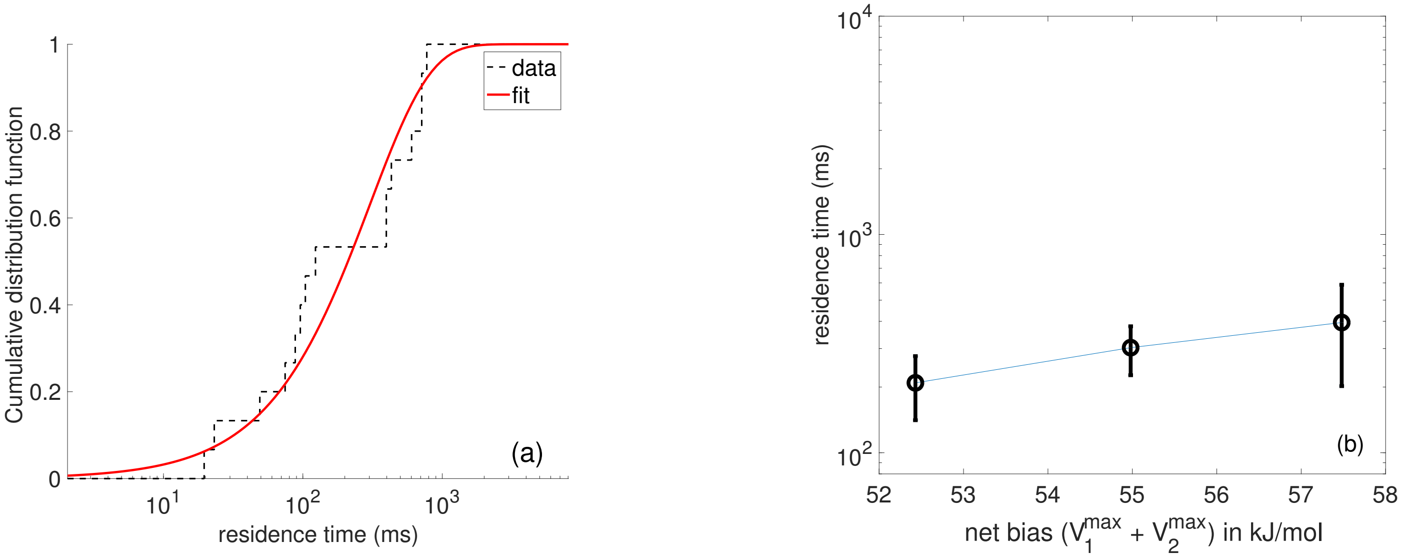


**Fig. S1.** (a) Poisson fit as per Ref.(5) to the 15 independent transition times obtained for the benzene-lysozyme dissociation constant with net maximum bias $V_1^{max} + V_2^{max}$ = 52.43 kJ/mol. The $k_{off}$ reported in the main text is calculated as the inverse of the fitted time constant here, $303 \pm 76$ ms. In (b), we provide the fitted time constant with associated error bars for 3 different biasing protocols, demonstrating convergence of the estimated time constant, at least on a log-scale. It is worth noting that for the weakest bias, we had the lowest number of dissociation events and hence the time-constant is a lower bound due to not having captured enough slow events. Hence the agreement should in principle be even better.

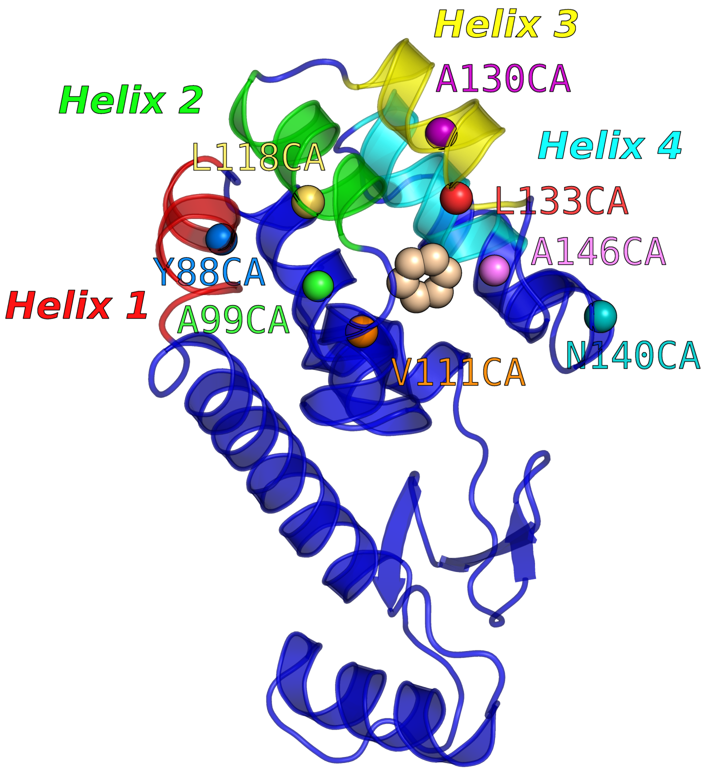


**Fig. S2.** Definition of Order parameters for Benzene-Lysozyme dissociation: Our 11 order parameters comprise 8 protein-ligand contacts and 3 protein-protein contacts. The protein-ligand contacts are implemented through the distances between the centres-of-mass of ligand to different protein atoms labeled in the plot. The protein-protein contacts are implemented through the distance between the centres-of-mass of ligand to centres-of-mass of helices labeled in the plot. Further details are provided in Table S1.

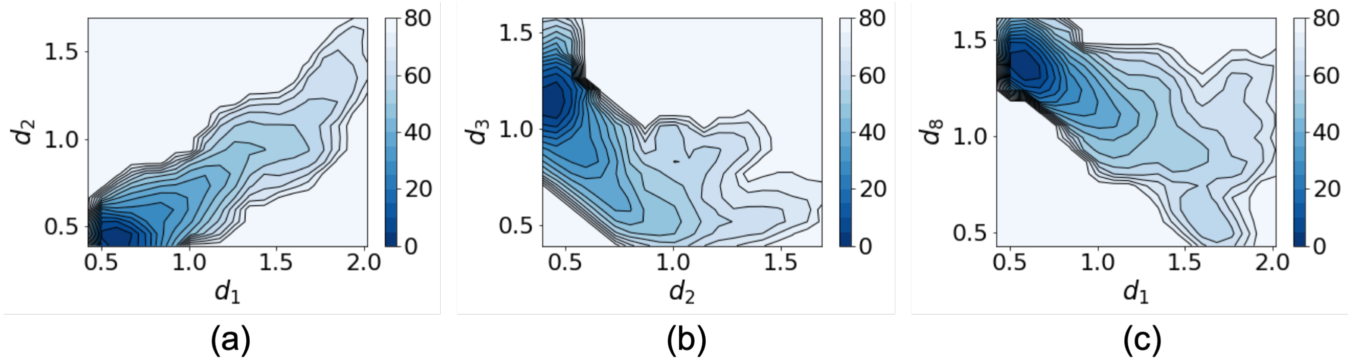


**Fig. S3.** Representative selected free energy surfaces for Benzene-Lysozyme dissociation obtained from our approach: (a) Free energy along $d_1$ and $d_2$. (b) Free energy along $d_2$ and $d_3$. (c) Free energy along $d_1$ and $d_8$. All energies are in units of kJ/mol with contours every 5 kJ/mol. These surfaces are in excellent agreement with previous benchmarks on this system (4).

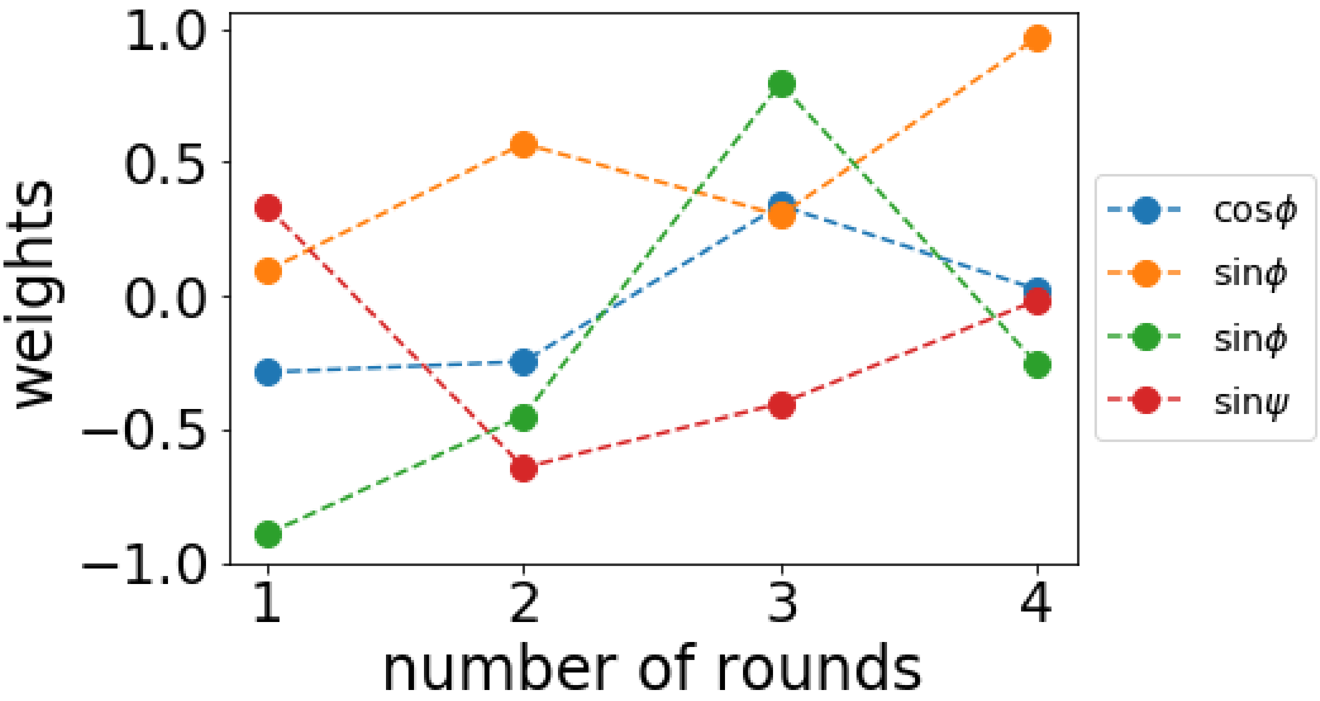


**Fig. S4.** Alanine dipeptide in vacuum: The value of weights for 4 order parameters a function of number of rounds.

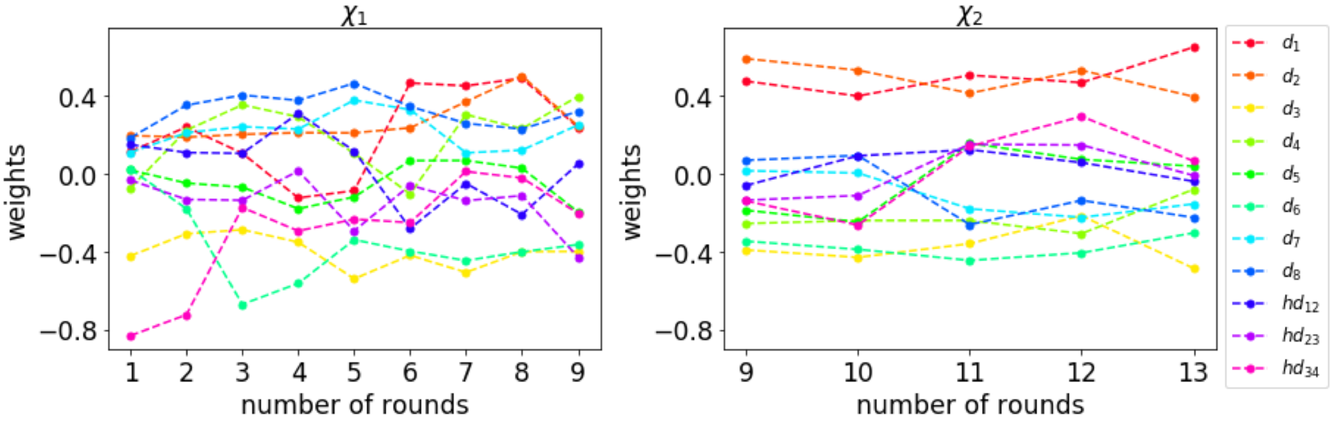


**Fig. S5.** Benzene-lysozyme dissociation: The plot on left shows the value of weights for 11 order parameters in $\chi_1$ as a function of number of rounds. The plot on right shows the value of weights for 11 order parameters in $\chi_2$ as a function of number of rounds. The plot on the right begins at 9 rounds directly because $\chi_2$ was not trained until 8 rounds.

| **Order parameter** | **Type** | **Definition** | **Weight $c_i$ in optimized $\chi_1 = \sum c_i s_i$** | **Weight in optimized $\chi_2 = \sum c_i s_i$** |
|---|---|---|---|---|
| $d_1$ | Protein-ligand | Y88CA–ligand | 0.4930 | 0.6524 |
| $d_2$ | Protein-ligand | A99CA–ligand | 0.5059 | 0.3984 |
| $d_3$ | Protein-ligand | L133CA–ligand | -0.4002 | -0.4853 |
| $d_4$ | Protein-ligand | L118CA–ligand | 0.2335 | -0.0792 |
| $d_5$ | Protein-ligand | V111CA–ligand | 0.0300 | 0.0399 |
| $d_6$ | Protein-ligand | A130CA–ligand | -0.4009 | -0.3037 |
| $d_7$ | Protein-ligand | N140CA–ligand | 0.1229 | -0.1544 |
| $d_8$ | Protein-ligand | A146CA–ligand | 0.2313 | -0.2240 |
| $hd_{12}$ | Protein-protein | Helix 1 (A82-S90) – Helix 2 (T115-123Q) | -0.2081 | -0.0378 |
| $hd_{23}$ | Protein-protein | Helix 2 (T115-123Q) – Helix 3 (W126-A134) | -0.1114 | -0.0073 |
| $hd_{34}$ | Protein-protein | Helix 3 (W126-A134) – Helix 4 (K147-T155) | -0.0204 | 0.0670 |

**Table S1. List of order parameters used to construct RC and their weights in $\chi_1$ and $\chi_2$.**

### References

1. Tiwary P, Parrinello M (2013) From metadynamics to dynamics. *Phys. Rev. Lett.* 111(23):230602–230606.
2. MacKay DJ, Mac Kay DJ (2003) *Information theory, inference and learning algorithms.* (Cambridge university press).
3. Valsson O, Parrinello M (2014) Variational approach to enhanced sampling and free energy calculations. *Phys. Rev. Lett.* 113(9):090601–090605.
4. Ribeiro JML, Tiwary P (2019) Towards achieving efficient and accurate ligand-protein unbinding with deep learning and molecular dynamics through rave. *Journal of chemical theory and computation.*
5. Salvalaglio M, Tiwary P, Parrinello M (2014) Assessing the reliability of the dynamics reconstructed from metadynamics. *J. Chem. Theor. Comp.* 10(4):1420–1425.
